# Rapid Parallel Visual Presentation Provides a New Perspective on Relative Clause Processing in Mandarin Chinese

**DOI:** 10.1101/2024.08.13.607840

**Authors:** Donald Dunagan, Jill McLendon, Tyson Jordan, Dustin A. Chacón

## Abstract

A fundamental question in language processing research is the degree to which the acceptability and ease of interpretation of a sentence are affected by the moment-by-moment memory encoding and retrieval processes needed to process it in time, vs. the properties of the linguistic representation itself, such as syntactic complexity. This manuscript investigates relative clauses, a crucial testing ground for theories of language processing, using parallel presentation, a technique in which the key elements of the target syntactic dependency are displayed to the visual system at the same time. Compared to serial presentation, parallel presentation mitigates the extent to which complex memory operations must be deployed to process the structure word-by-word. We report on an electroencephalography (EEG) experiment in Mandarin Chinese. Participants were presented with sentences which contained either a subject- or object-modifying subject- or object-extracted relative clause. Sentence presentation was split across two separate screens, with each screen being displayed only for 166ms on average. We find a behavioral preference for subject-extracted relative clauses and distinct EEG signatures for subject- and object-extracted relative clauses. Together, these results – on account of the parallel presentation scheme applied here – provide support for the independent contribution of linguistic representation in processing difficulty and the perspective that relative clause processing involves more than the factors that word by word accounts of this phenomenon would suggest. Further, the EEG result contributes to a growing body of electrophysiology literature attempting to illuminate the processing mechanisms deployed in the context of parallel presentation.

---

A fundamental question in language processing research is the degree to which the acceptability and ease of interpretation of a sentence are affected by the moment-by-moment memory encoding and retrieval processes needed to process it in time, vs. the properties of the linguistic representation itself, such as syntactic complexity. Appeals to representational complexity are less frequently invoked in psycholinguistic theory and are largely associated with the Derivational Theory of Complexity (Fodor & Garrett, 1967; but see Hofmeister & Sag, 2010). This manuscript investigates the processing of relative clauses, a crucial testing ground for theories of language processing difficulty (King & Just, 1991). However, we depart from prior research by using parallel presentation, in which the key elements of the target syntactic dependency are displayed to the visual system at the same time. This mitigates the extent to which complex memory operations must be deployed to process the structure word-by-word over time (Chacón et al., 2024; Krogh & Pylkkänen, 2024). Our research question, broadly, is: Is there still a detectable difference between subject- and object-extracted relative clauses when the entire syntactic dependency is displayed in parallel? If so, this would provide support for the role of structural representation in processing difficulty, apart from the factors suggested by word by word accounts. Here, we present on an electroencephalography (EEG) experiment in Mandarin Chinese, a language in which many conflicting findings have previously been reported (Yun et al., 2015).

Relative clauses are sentence-like units which modify a noun. In (1), the noun *boy* has been relativized (extracted) from its position as subject of the relative clause verb *loves*. This relativization creates a filler-gap relation (dependency), as indicated via the coreferentiality index

*i.* Relative clauses (RCs) can either be subject-extracted (1, 3) or object-extracted (2, 4), and can either be subject-modifying (1, 2) or object-modifying (3, 4) – among other possibilities not discussed here.

1. The [boy]*_i_* that *_i_* loves the dog sees the cat
2. The [dog]*_i_* that the boy loves *_i_* sees the cat
3. The cat sees the [boy]*_i_* that *_i_* loves the dog
4. The cat sees the [dog]*_i_*that the boy loves *_i_*

English has Subject Verb Object (SVO) word order and uses head-initial RCs; that is, the head (relativized) noun occurs before the RC which modifies it (alternatively, RCs are *postnominal* in English). In contrast, Mandarin Chinese (from here forward simply *Chinese*), which is also SVO, uses *head-final* RCs; that is, the head noun occurs after the relative clause which modifies it (alternatively, RCs are *prenominal* in Chinese). It is not the case, however, that all languages with head-final RCs are SVO. Japanese and Korean, for example, use head-final RCs, but have Subject Object Verb (SOV) word order. As can be seen in (5-8), Chinese RCs are characterized by the relativizer DE (的), which precedes the relativized head noun. It is because of the different realizations across languages that relative clauses provide a fertile testing ground for the investigation of human sentence processing.

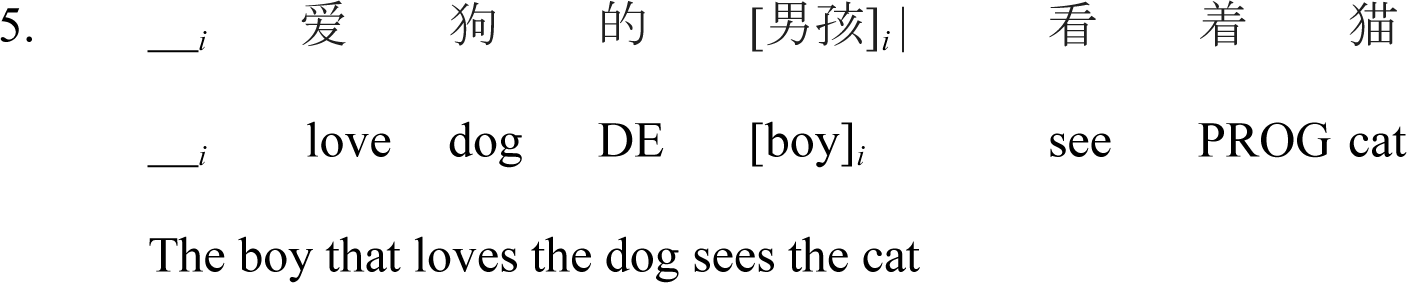

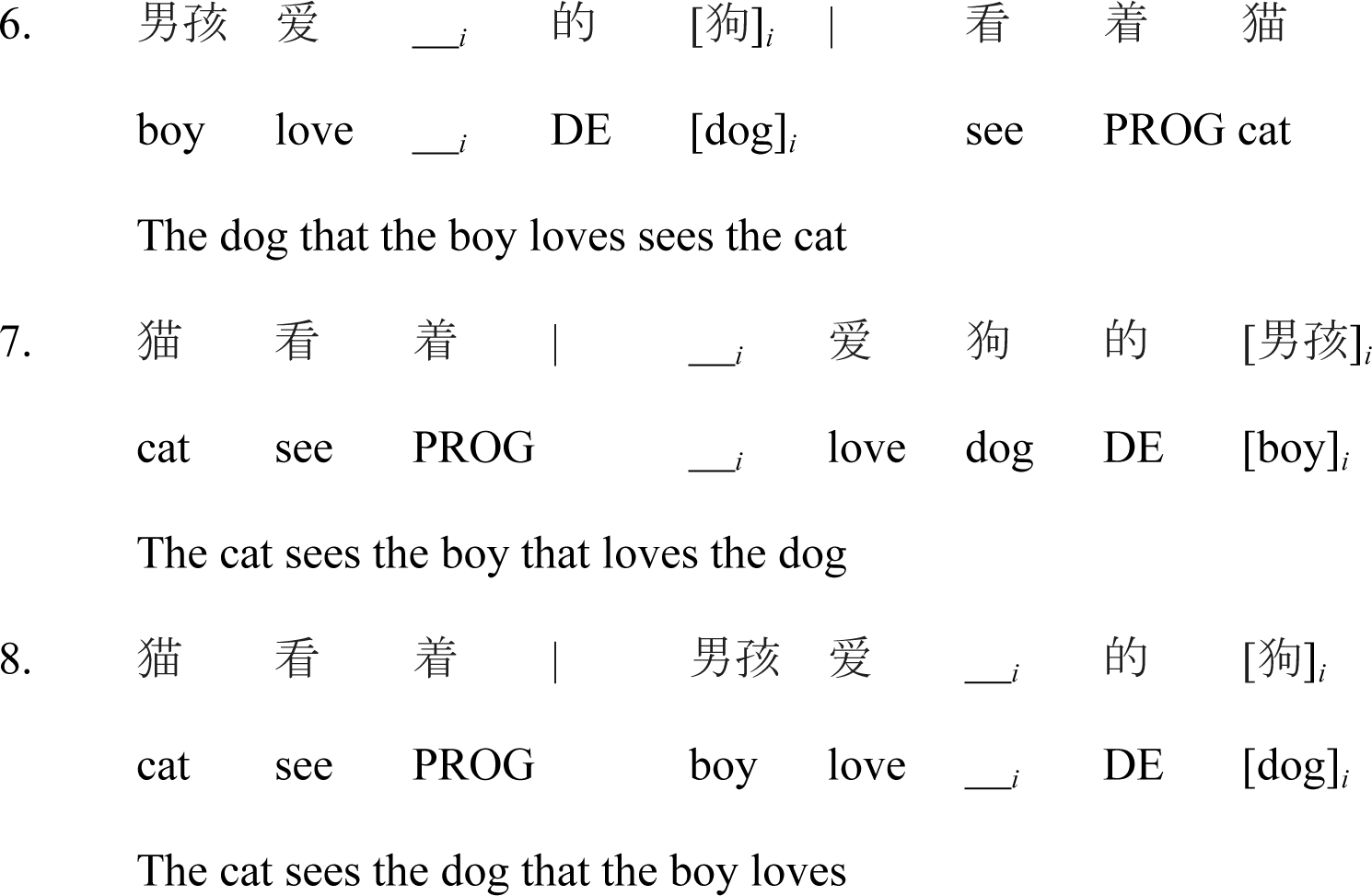

It is well-documented in the sentence processing literature that, in English, subject-extracted relative clauses (SRCs) are easier to process than object-extracted relative clauses (ORCs); that is, there exists an SRC *advantage* (see Lau & Tanaka, 2021 for a comprehensive overview of RC processing asymmetry). Evidence converges from reading times (Gibson et al., 2005; King & Just, 1991; Staub, 2010; Traxler et al., 2002, 2005), question answering (Wanner & Maratsos, 1978), event related potentials (ERPs; King & Kutas, 1995; Müller et al., 1997), and neuroimaging (Caplan et al., 2008; Just et al., 1996; Stromswold et al., 1996). While previous work has found an SRC advantage, like in English, for Korean (Kwon et al., 2006, 2010, 2013; O’Grady et al., 2003) and Japanese (Ishizuka, 2005; Miyamoto & Nakamura, 2013; Ueno & Garnsey, 2008) – which use head-final RCs and SOV word order – evidence is more mixed for Chinese. Some reading (Chen et al., 2008; Gibson & Wu, 2013; Hsiao & Gibson, 2003; Lin, 2014; Lin & Garnsey, 2010; Sung, Cha et al., 2016; Sung, Tu et al., 2016; Xu et al., 2019; Xu et al., 2020a), ERP (Packard et al., 2011; Xiaoxia et al., 2016), and neuroimaging (Xu & Duann, 2020; Xu et al., 2020a, 2020b) studies have found an ORC advantage – the opposite of English, Korean, and Japanese. However, other reading (Jäger et al., 2015; Li et al., 2010; Lin & Bever, 2006, 2010; Vasishth et al., 2013), ERP (Xiong et al., 2019), and information-theoretical (Chen et al., 2012; Yun et al., 2015) analyses have found an SRC advantage – as in English, Korean, and Japanese. To complicate matters further, some Chinese studies have found either no advantage (Lee & Chan, 2023, fMRI; Zhou et al., 2018, reading time), that the advantage changes depending upon location in the RC (Bulut et al., 2018, ERP; Wang et al., 2017; Yang et al., 2010), or that the advantage changes depending on whether the RC is subject- or object-modifying (Xiong & Newman, 2021, fMRI).

This paper aims to bring a new perspective to the Chinese RC processing discussion by applying a relatively new experimental paradigm: rapid parallel visual presentation. In contrast to the common experimental paradigm of presenting participants with one word at a time (rapid serial visual presentation; RSVP), recent work (Asano & Yokosawa, 2011; Chacón et al., 2024; Declerck et al., 2020; Defau et al., 2024; Dunagan et al., 2024; Fallon & Pylkkänen, 2023; Flower & Pylkkänen, 2024; Krogh & Pylkkänen, 2024; Massol et al., 2021; Snell & Grainger, 2017) has investigated language processing in the context of visually presenting participants with short, sentence-length stimuli, but only briefly – around 200-300ms (rapid parallel visual presentation; RPVP). From this direction of investigation, a *sentence superiority effect* (SSE) emerges such that participants perform better in behavioral tasks in response to grammatical and semantically congruent stimuli than ungrammatical and semantically anomalous stimuli, suggesting that some grammatical processing can occur at this presentation time. Recording EEG, Wen et al. (2019; 2021a) observe a reduced N400 for grammatical compared to scrambled sentences. This EEG SSE for grammatical vs. scrambled conditions was replicated by Dunagan et al. (2024), who further found that in the RPVP paradigm, the human sentence processor is neither behaviorally nor electrophysiologically sensitive to subject-verb number violation.

Chacón et al. (2024) found that sentences with *wh*-constructions elicited distinct neural signatures from their non-*wh* counterparts 300–400ms post-sentence onset, for both *wh*-filler-gap dependencies (like English) and *in-situ wh*-phrases, much earlier than the usual P600 reported for filler-gap dependencies, lending further support to the claim that RPVP may mitigate the demands on working memory imposed by serial reading experiment designs.

Some of the recent work with RPVP has made use of magnetoencephalography. Using the same stimuli and grammatical vs. scrambled contrast as Snell & Grainger (2017) and Wen et al. (2019), Dufau et al. (2024) identify a sequence of left-lateralized language network regions as being associated with the SSE: first the inferior frontal gyrus (321-406ms), then the anterior temporal lobe (466-531ms), and lastly the inferior frontal (549-602ms) and the posterior superior temporal gyri (553-622ms). Investigating the *transposed-word effect* (Dufour et al., 2022; Pegado et al., 2021; Wen et al., 2021a, 2021b, 2022) – a phenomenon in which ungrammatical stimuli that contain two words that can be transposed to create a grammatical construction are privileged over ungrammatical stimuli which cannot be made grammatical via transposition – Flower & Pylkkänen (2024) discover that the ungrammaticality of transposed stimuli (*all are cats nice*) is quickly noticed (213ms), but then rapidly ‘fixed’ (468ms). Krogh & Pylkkänen (2024) use RPVP – in the context of syntactic dependency, syntactic frame, and verb argument structure in Danish 2-word sentences – to probe the neural correlates of syntax, unbound from the working memory burden necessarily imposed by traditional serial presentation.

Many theories abound proposing to account for (cross-linguistic) RC processing asymmetry. Under the Parallel Function Hypothesis (PFH; Sheldon, 1974), RC processing is facilitated when the coreferential matrix and relative noun phrases have the same grammatical function (i.e., subject-modifying SRCs and object-modifying ORCs). The Perspective Hypothesis (MacWhinney, 1977, 1982; MacWhinney & Pléh, 1988) affords easier processing to constructions which maintain rather than shift perspective. Some models (Chen & Hale 2021; Chen et al., 2012; Yun et al., 2015) make use of information-theoretic complexity metrics (e.g., Hale, 2001, 2003, 2006, see 2016 for a review; Levy 2008) to derive processing asymmetry. Lin (2014, 2015) posits that RCs with canonical thematic order are easier to process. The Accessibility Hierarchy (AH; Keenan & Comrie, 1977; Keenan & Hawking, 1987) develops a hierarchy of relativization positions. Positions higher up in the hierarchy are more accessible to relativization, and relativizations created from higher up positions are easier to process. Under Gibson’s Dependency Locality Theory (DLT; 2000), difficulty results from two separate components: having to store more unresolved syntactic dependencies in working memory, and having to integrate incoming words into the current structural representation over a larger distance. DLT predicts an SRC advantage in languages with head-initial RCs like English, but an ORC advantage in languages with head-final RCs like Korean, Japanese, and Chinese, since these relativization dependencies cross fewer words. In contrast, under the Structural Distance Hypothesis (SDH; O’Grady, 1997; O’Grady et al., 2003) processing difficulty is determined not by *linear* distance, but by *structural* distance between the extracted element and its originating position. Distance is calculated, for example, in terms of number of intervening syntactic nodes between the filler and the gap in the syntactic representation. As objects will always be more deeply embedded (originate lower in the phrase structure) than subjects, the SDH predicts a cross-linguistically consistent SRC advantage, regardless of RC head directionality or the number of intervening words.

Accounts of relative clause processing can (roughly) be divided based upon whether there is a role for temporality. While some proposals – like accounts which build off of word-by-word disambiguation and the Perspective Hypothesis – necessarily consider time and word-by-word memory operations, others – like the AH and the SDH – are time agnostic, and instead attribute the differences between SRCs and ORCs to differences in their time-independent representations. The RPVP presentation scheme applied here allows for valuable insight into the psycholinguistic correlates of grammatical structure processing as it *abstracts* over temporality, temporal ambiguity, and working memory demands (Chacón et al., 2024; Krogh & Pylkkänen, 2024).

Here, we investigate RC processing in Chinese using RPVP and EEG. In English, ORCs, relative to SRCs, have been shown to elicit a (sustained) anterior negativity (SAN) between the filler and the gap site, with the interpretation being that this is a reflection of increased working memory demand (King & Kutas, 1995; Müller et al., 1997): in English ORCs, the matrix subject must be maintained in working memory during the processing of the RC. In Chinese, however, event-related potential (ERP) correlates of RC processing are quite varied (in line with the diversity of the broader collection of SRC vs. ORC reports cited above). Packard et al. (2010) investigate the P600 – a positive-going ERP component that correlates with processing difficulty and computing more complex grammatical representations. For subject-modifying RCs, they find an increased P600 at the relativizer DE for SRCs; for object-modifying RCs, they find an increased P600 for SRCs at the RC head; they interpret their P600 results as reflecting increased syntactic integration difficulty for the SRCs. Looking at object-modifying RCs, Yang et al. (2010) identify: a P600 for SRCs at the RC verb – which they interpret as *structural* reanalysis, an N400 for ORCs at the at the RC verb – which they interpret as *meaning* reanalysis, and a sustained negativity for ORCs at the head noun – which they interpret as comprehension and referent resolution. At the relativizer, Bulut et al. (2018) identify both an N400 – driven by their low working-memory group – and a P600 for ORCs. They interpret this complex effect as resulting from the detection of ambiguity and subsequent reanalysis. At the head noun, they identify a P600 for SRCs, which they interpret as reflecting syntactic integration difficulty. At the relativizer, Xiong et al. (2019) identify an early left anterior negativity (eLAN) for ORCs – which they interpret as increased working memory demand during gap-filler integration, and for object-modification, a P600 for ORCs – which they interpret as reflecting structural reanalysis.

This paper contributes: a) to the Chinese RC processing (EEG) literature by investigating relative clause processing using RPVP; and b) to the blossoming RPVP paradigm by presenting a novel adaptation to this protocol in which participants are presented with two, back-to-back, rapidly presented screens. In this manner, stimuli that would otherwise be too long to process in a single glance (pertinently Chinese sentences with relative clauses) can be investigated using RPVP. RPVP creates an intriguing opportunity for the case of Chinese relative clauses because it removes the temporal component associated with word-by-word processing; temporality is a complicating factor for Chinese RC processing, both because of the long-distance dependency and local ambiguities. Because Mandarin Chinese allows for null pronouns, comprehenders may not immediately detect gaps, and because the relativizer follows the relative clause, comprehenders may not consider a relative clause parse until after encountering 的 DE (e.g., Hsiao & Gibson, 2003; Lin & Bever, 2010). Further, removing the temporal component neutralizes many prominent theories of RC processing, including DLT and word-by-word disambiguation. Our specific research questions are, in the context of Chinese RCs presented in RPVP: a) Whether an SRC or ORC advantage is observed; and b) whether any traditionally-observed RC-associated ERPs (i.e., (e)LAN, N400, P600, SAN) are observed.

## Method

### Participants

Thirty-four self-identified Mandarin Chinese speakers were recruited from the Athens, Georgia community. One participant was removed due to experimenter error and three participants were removed due to poor performance on the task (< 70% accuracy). Each participant’s language history was documented using the Mandarin Chinese version of the Leap-Q questionnaire (Marian et al., 2007). Participants were also interviewed about their reading experience in Chinese, to ensure that they had substantial facility in reading Chinese script. Handedness was also tracked using the Edinburgh Handedness Survey (Oldfield, 1971). Post-hoc analyses revealed no meaningful relation between handedness, language history, and task performance, and we did not detect any systematic differences between participants’ recorded EEG signals that could be explained by any of the demographic variables. Participants all gave written informed consent before participating, and were provided with consent forms in Chinese, or another language by request. Initial recruitment communications occurred in the target languages by a native speaker experimenter.

### Materials

A total of 48 sets of sentences were prepared in 4 conditions, distributed in a 2 × 2 design Each set consisted of simple SVO sentence with a transitive verb and a single relative clause.We manipulated whether the RC modified the subject NP or object NP (±Object-Modifying), and whether the RC was a subject-extracted relative clause or an object-extracted relative clause (±ORC). All nouns were plausible agents: either an animal or a human agent. An example stimulus set is given in (5-8) and Figure 1B.

**Figure 1.**
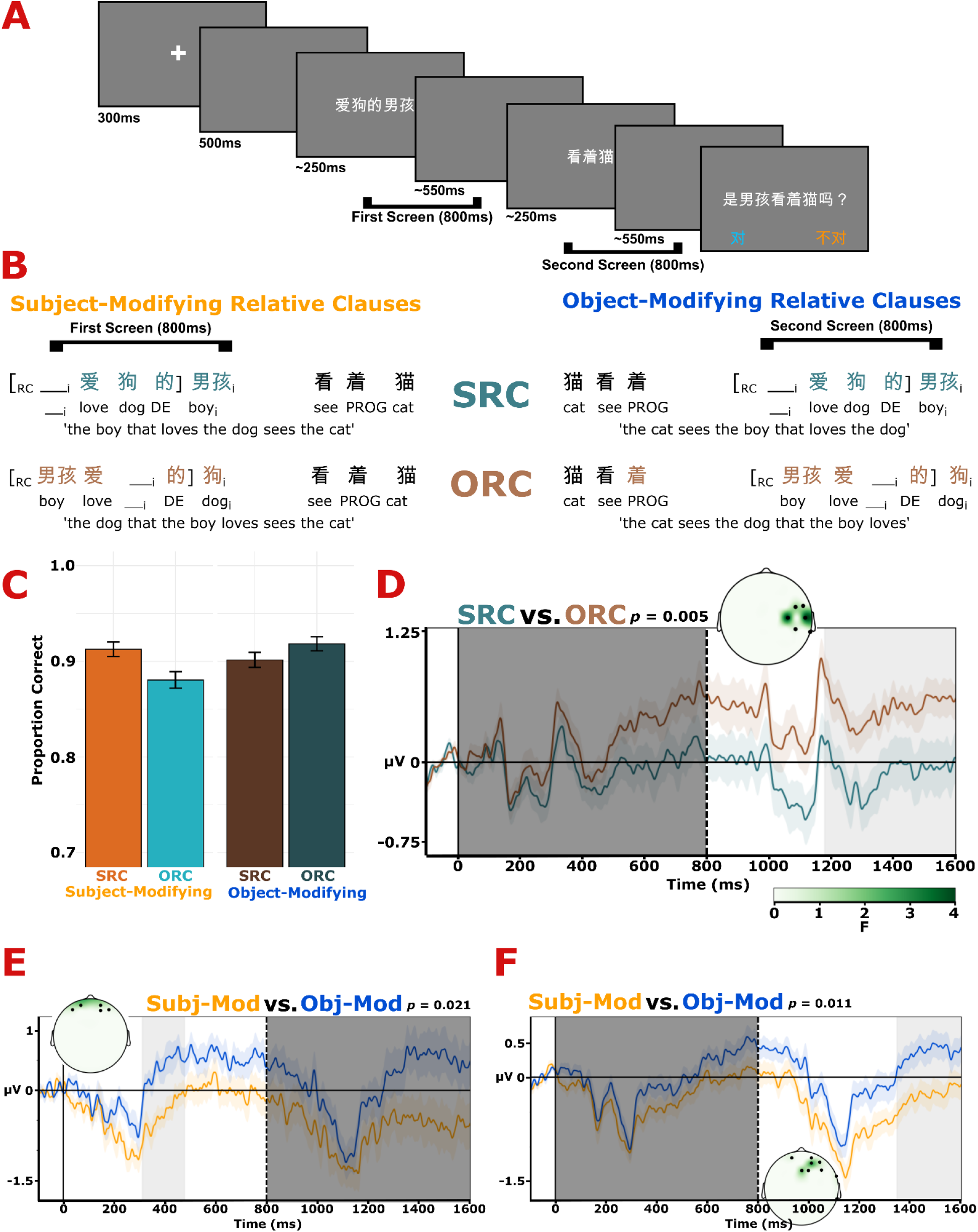
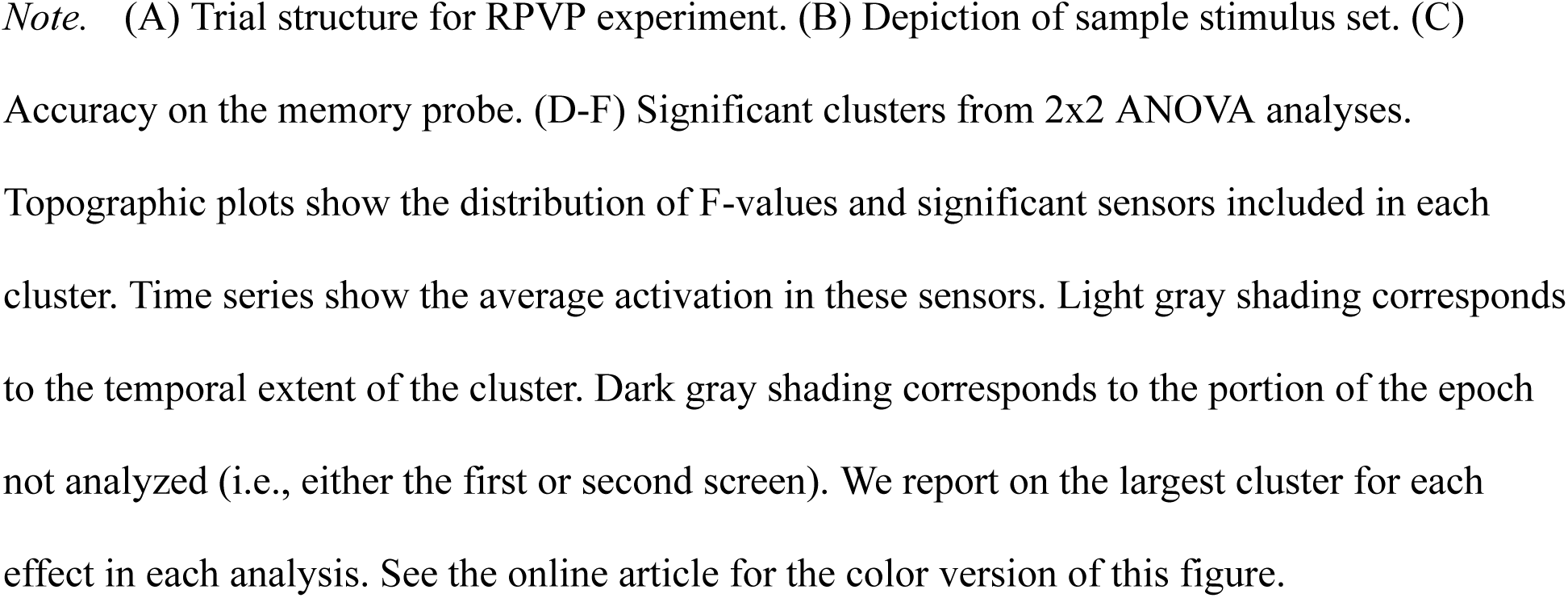
Experiment Structure, Sample Stimulus Set, and Behavioral and EEG Results.

### Procedure

The rapid parallel visual presentation experiment was conducted using PsychoPy. The procedure builds off of previous RPVP schemes (e.g., Asano & Yokosawa, 2011; Chacón et al., 2024; Flower & Pylkkänen, 2024; Snell & Grainger, 2017). Stimuli were presented within-subjects, such that each participant saw each sentence. Order presentation was pseudo-randomized into 4 separate lists, such that one item set occurred in one condition per list, and each list contained an equal number of trials per condition. Between each list, participants took a break, in which participants saw photos of baby animals.

Each sentence was displayed in two halves after a fixation cross. This was done because it was not feasible to have the entire Chinese sentence viewable within the fovea and parafovea. We ensured that the entire relative clause and its head were visible on the same screen. For the subject-modifying RC conditions, this means that subject NP occurred on the first screen, and the main verb and object NP occurred on the second. For the object-modifying RC conditions, this means that the subject NP and the main verb occurred on the first screen, and the object NP occurred on the second.

Each trial was preceded by a fixation cross, centered in the screen and displayed for 300ms, followed by a blank screen for 500ms. Then, the sentence was displayed in two screens. Each screen appeared for a fixed duration of time, which varied between 150ms – 600ms, and was followed by a blank screen. The duration of the screens with text and the blank screens were dynamically adjusted according to participants’ performance, and were constrained such that each text fragment-blank screen pair lasted for 800ms. Finally, an untimed comprehension question occurred after the second blank screen. Participants entered their response by pressing the [F] key to indicate ‘true’, and the [J] key to indicate ‘false.’ On-screen reminders were provided on the lower half of the screen during the comprehension question. An error message that lasted for 1000ms was provided as on-screen feedback for incorrect responses. The trial structure is exemplified in Fig. 1A. We elected to use comprehension questions instead of memory probes (Snell & Grainger, 2017; Wen et al., 2019, 2021) or a sentence match/mismatch task (Dunagan et al., 2024; Flower & Pylkkänen, 2024; Krogh & Pylkkänen, 2024) because it was crucial that comprehenders assigned the correct thematic relations to the argument NPs in the sentence. Following Dunagan et al (2024) and Chacón et al. (2024), the presentation times of the two stimulus screens were dynamically adjusted based upon participant performance on the comprehension questions. We did this because, to our knowledge, complex sentences have not yet been investigated with a RPVP design. Previous findings find sentence superiority effects with presentation times 200–300ms (Dunagan et al., 2024; Flower & Pylkkänen, 2024; Krogh & Pylkkänen, 2024; Snell & Grainger, 2017; Wen et al., 2019, 2021), suggesting that some syntactic information can be gleaned at this presentation rate. However, our stimuli are presented with Chinese characters and our stimuli are multiclausal, and may therefore require extra processing time, comparatively. Presentation times were dynamically adjusted such that incorrect answers resulted in the stimuli presented for an additional 50ms per screen, with 50ms deducted from the blank screen time. Similarly, correct answers resulted in a reduction of 50ms per screen, with 50ms added to the blank screen time.

Participants were kept at a uniform distance from the screen, approximately 70cm from nasion to center of the screen. The visual angle subtended of the stimuli was approximately 15 degrees. All stimuli were printed in a white font (ST Heiti Light) against a dark gray background in a dimly-lit room. All on-screen instructions and communications were conducted in Chinese.

EEG signals were recorded using a 64 channel Ag/Cl BrainVision actiChamp+ system (Gilching, Germany). Impedance of the EEG sensors was reduced by the application of SuperVisc gel, and lowered to <20kΩ. On-line EEG recording was referenced to FCz according to manufacturer standards, and then re-referenced to an average reference offline.

Participants engaged in an unrelated task that is not reported here. The order of the two tasks was counterbalanced. Participants also engaged in a series of localizer tasks at the beginning of each recording session, but we do not report this here.

### Transparency and Openness

All materials, data, stimuli, and preparation, presentation, and analysis code are available at https://osf.io/ub3x9/. The design and analysis of this study were not preregistered.

## Results

### Behavioral Results

We used the lme4 package (Version 1.1-35.1) in R (Version 4.3.2,) to fit a logistic mixed effects regression model to the accuracy data. We fit the correct response as the dependent variable, with ±Obj-Modifying, ±ORC, and their interaction as predictors, and with participant and item as random effects. More complex random effects structures failed to converge. The model was fit with Subject-Modifying and Subject-Extracted coded as 0 and Object-Modifying and Object-Extracted coded as 1. The results of the model are shown in Table 1, and the average accuracy by condition is shown in Figure 1C.

**Table 1.** Results of Logistic Mixed Effects Model fit to the Accuracy Data.

| | $\beta$ | SE | $z$ | $p$ |
| --- | --- | --- | --- | --- |
| *(Intercept)* | 2.61 | 0.17 | 15.63 | **< .001** |
| *Obj-Mod* | -.14 | 0.13 | -1.08 | .28 |
| *ORC* | -.37 | 0.13 | 2.97 | **< .01** |
| *Obj-Mod x ORC* | .59 | 0.18 | 3.23 | **< .01** |

*Note.* The structure of the model was Correct $\sim \pm$ Obj-Modifying $\ast \pm$ ORC + (1|Participant) + (1|Item). The model was fit with Subject-Modifying and Subject-Extracted coded as 0 and Object-Modifying and Object-Extracted coded as 1. Bolded $p$ -values are significant at alpha < .05.

Overall, participants did very well on the task (mean accuracy 90.31%, SE 0.004). Because participants performed near-ceiling on the behavioral task, the presentation times were overwhelmingly low, concomitantly, with an average presentation time of 166ms (SE = 0.7ms) per screen.The mixed effects logistic regression on comprehension accuracy indicates that while there is no difference in performance for subject-vs. object-modification (*p* > .05), participants perform significantly worse on sentences with ORCs compared to those with SRCs (*p* < .01). Notably, a significant interaction was found such that ORC processing was facilitated in the object-modifying condition (*p* < .01).

### EEG Analysis

All EEG processing was conducted in MNE-Python (Version 1.4, Gramfort et al., 2013), and statistical analysis was conducted in Eelbrain (Version 0.39.5, Brodbeck et al., 2023). Raw EEG data were recorded using FPz as reference, following manufacturer instructions. Raw EEG data were then band-pass filtered off-line using an IIR Butterworth filter from 0.1–40Hz. Flat and noisy channels were then identified, removed, and interpolated. We then re-referenced EEG data to an average reference. Then, independent components analysis was used to identify and remove endogenous sources of rhythmic electromagnetic noise, including eye-blinks, ocular movements, and heartbeats. Afterwards, epochs were extracted for the target sentence, from 100ms pre-stimulus onset to 1600ms post-stimulus onset, encompassing both halves of the stimuli. Incorrect trials were excluded at this stage. Epochs were baseline corrected to the –100-0ms pre-onset period. Epochs with extreme voltage fluctuations were automatically rejected with a 100μV peak-to-peak threshold, and other bad epochs were rejected after visual inspection. For statistical analyses, condition numbers were normalized by subject, and then condition averages were computed. After rejection of incorrect trials and bad epochs, and by-subject condition normalization, 95.2% of trials remained for analysis.

After pre-processing, the sensor data was analyzed using spatio-temporal cluster-based permutation tests (Maris & Oostenveld, 2007). This method allows us to leverage the fact that adjacent time points and sensors are not independent to identify significant clusters in space-time, without the problem of implicit multiple comparisons used in pre-selecting sensors and time windows of analysis. Two analyses were performed: one in the time window from 200–800ms (first screen) and one in the time window from 1000–1600ms (second screen). For each time point and sensor, we fit an independent ANOVA to the raw EEG activity across individuals with ±Obj-Mod and ±ORC as factors (μV ∼ ±Obj-Mod * ±ORC). –Obj-Modifying and –ORC were coded as 0 and +Obj-Modifying and +ORC coded as 1. This produces an F-value test-statistic and a *p*-value for each time point and sensor. Afterwards, points are clustered together if they are adjacent in time and space. This clustering procedure is constrained to clusters of minimum length 20ms, minimum 3 sensors, with contributing points *p* < .05. All EEG sensors were included in the spatio-temporal clustering procedure. Afterwards, the test-statistics (F-values) were summed to produce the cluster-level statistic (’cluster size’). The ANOVA and clustering procedure was then conducted another 10,000 times, each time randomly permuting the condition labels on the data. These clusters are then ordered by size. This produces a null distribution against which to compare the clusters identified in the observed data. The observed data’s cluster’s *p*-values after correction are their ranks in the bootstrapped null distribution; clusters in the top 5% of this distribution are considered significant at α < .05.

## EEG Results

### First Screen

For the first screen two clusters were observed for ±Obj-Mod. The largest was an anterior cluster between 309–477ms with greater positivity for object-modifying RCs than subject-modifying RCs (*p* = .021; Figure 1E). The smaller cluster occurred over left lateral electrodes between 331-413ms with greater positivity for subject-modifying RCs than object-modifying RCs (*p* = .045). We report on the larger cluster only in Figure 1. No significant clusters were observed for ±ORC, nor were any significant clusters observed for the interaction of ±ORC and ±Obj-Mod.

### Second Screen

For the second screen, one cluster was observed for ±ORC over right lateral electrodes between 1179–1599ms (379–799ms post second screen presentation) with greater positivity for ORCs than SRCs (*p* < .01; Figure 1D). Two clusters were observed for ±Obj-Mod. The larger ±Obj-Mod cluster occurred over right anterior electrodes between 1345–1600ms (545–800ms post second screen presentation) with greater positivity for object-modifying RCs than subject-modifying RCs (*p* = .011; Figure 1F). The smaller ±Obj-Mod cluster also occurred over right anterior electrodes between 1000–1255ms (200–455ms post second screen presentation), but with greater negativity for subject-modifying RCs than object-modifying RCs (*p* = .014). We report on the larger cluster only in Fig. 1. No significant clusters were observed for the interaction of ±ORC and ±Obj-Mod.

## Discussion

Our research questions were: a) whether an SRC or ORC advantage would be observed for Chinese RCs presented in RPVP; and b) if found, whether an observed difference could be associated with traditionally-observed RC-associated ERPs. With respect to the first, the behavioral results to the comprehension questions indicate an SRC advantage where participants perform worse on comprehension questions from ORC stimuli than they do on questions from SRC stimuli. Further support for an SRC advantage comes from the EEG data, in which, during the time window of the second screen, ORCs were associated with greater right lateral positivity. ORC vs. SRC processing, then – on account of the parallel presentation scheme applied here – involves more than the factors that word-by-word accounts that involve working memory would suggest. To this end, we suggest that the observed SRC advantage results from the structural difference between SRCs and ORCs (Keenan & Comrie, 1977; Keenan & Hawking, 1987; O’Grady, 1997; O’Grady et al., 2003). Interestingly, we also observe an interaction effect in the behavioral data in which ORC processing is facilitated when the ORCs are object-modifying, suggesting that RC processing is in part shaped by the role that the head noun plays in the main clause (cf. Sheldon, 1974). In regard to the second research question, the observed cluster distinguishing SRCs from ORCs does not resemble typical RC or long distance dependency-associated ERPs (i.e., (e)LAN, N400, P600, SAN). This is in line, however, with recent work by Chacón et al. (2024) who find that while *wh*-constructions presented in RPVP elicit distinct neural signatures from their non-*wh* counterparts, these signatures do not resemble the usual P600 reported for filler-gap dependencies in serial presentation.

### Subject-Modifying vs. Object-Modifying

For both screens, main effect clusters are observed in the EEG data for subject-vs. object-modification. This, however, is an expected artifact of the presentation scheme, where the subject and object of the sentence are split across two screens: for both screens, the lexical material and syntactic structure (the presence or absence of an RC) differ between the +Object-Modifying and –Object-Modifying conditions. Indeed, no main effect of subject-vs. object-modification was observed in the behavioral data. The largest clusters share similar frontal topographies, and in both, object-modification is associated with greater positivity and subject-modification with greater negativity. The earliest cluster of the first screen began around 300ms, while the earliest cluster of the second screen (which also exhibits a frontal topography) began around 200ms post second screen presentation. Previous work by Chacón et al. (2024) – investigating ±WH constructions in Chinese with single screen RPVP presentation – found three clusters indicating that WH object constructions were rapidly distinguished from NP object constructions as early as 200ms (∼200ms, ∼200ms, ∼300ms), with two of the clusters occurring with an anterior topography, like the clusters observed here. In this context, the results of this investigation provide further evidence to support the claim that the human sentence processor is able to rapidly process syntactically complex constructions when stimuli are presented in a parallel manner.

### Subject-Extracted Relative vs. Object-Extracted Relative

The behavioral results reveal a subject-vs. object-extracted relative clause asymmetry in which participants perform worse in the ORC condition than the SRC condition. This contrasts against previous results which have found an ORC advantage (e.g., Gibson & Wu, 2013; Hsiao & Gibson, 2003; Lin & Garnsey, 2010), and instead, is in line with results which have found an SRC advantage (e.g., Jäger et al., 2015; Lin & Bever, 2010; Vasishth et al., 2013). Further SRC advantage support is provided by the EEG data in which, following the second screen, greater positivity is found for ORCs than SRCs over right lateral electrodes ∼1200ms (∼400ms post second screen presentation), which may be interpreted as a P600 compatible with previous findings (Bulut et al., 2018; Xiong et al., 2019) To note, the onset of the RCs were not synchronized between the levels of ±Object-Modifying: subject-modifying RCs occurred with the subject NP in the first screen, and object-modifying RCs occurred with the object in the second screen. This means that the main effect of ±ORC observed in the EEG data should not be interpreted with respect to the onset of the RCs. However, the absence of an effect for the first screen and the absence of an interaction in either screen, combined with the presence of an effect for the second screen (the RC could have occurred in the first screen and not the second), indicate that the SRC vs. ORC processing contrast, either subject- or object-modifying, didn’t manifest until the entire sentence had been presented and was available for processing.

Of the accounts of RC processing which are compatible with RPVP, the integration aspect of DLT (Gibson, 2000) predicts an ORC advantage, the SDH (O’Grady, 1997; O’Grady et al., 2003) and AH (Keenan & Comrie, 1977; Keenan & Hawking, 1987) predict an SRC advantage, and the PFH predicts an advantage for subject-modifying SRCs and object-modifying ORCs. Proposals that necessarily involve serial processing will not be further considered here, in the context of RPVP. In DLT, integration difficulty is (partially) determined by the amount of linear material that occurs between a filler and its gap site. In Chinese, there is more intervening material for SRCs, than ORCs. In contrast, in the SDH – which can be thought of as an implementation of the AH – difficulty is thought to result from structural distance (i.e., number of syntactic nodes) between the filler and its gap site. Figure 2 illustrates syntactic trees for example Chinese subject- and object-extracted relative clauses, annotated with the amount of structural material that occurs between the filler and the gap site. As can be seen, there are a greater number of intervening syntactic nodes in the ORC (6; N, NP, RC, S, VP, NP) than the SRC (5; N, NP, RC, S, NP). Thus, greater processing/integration difficulty is predicted for ORCs compared to SRCs. Indeed, in observing both a behavioral and electrophysiological SRC advantage, the results of this study support the Structural Distance Hypothesis, as compared to Dependency Locality Theory.

**Figure 2.**
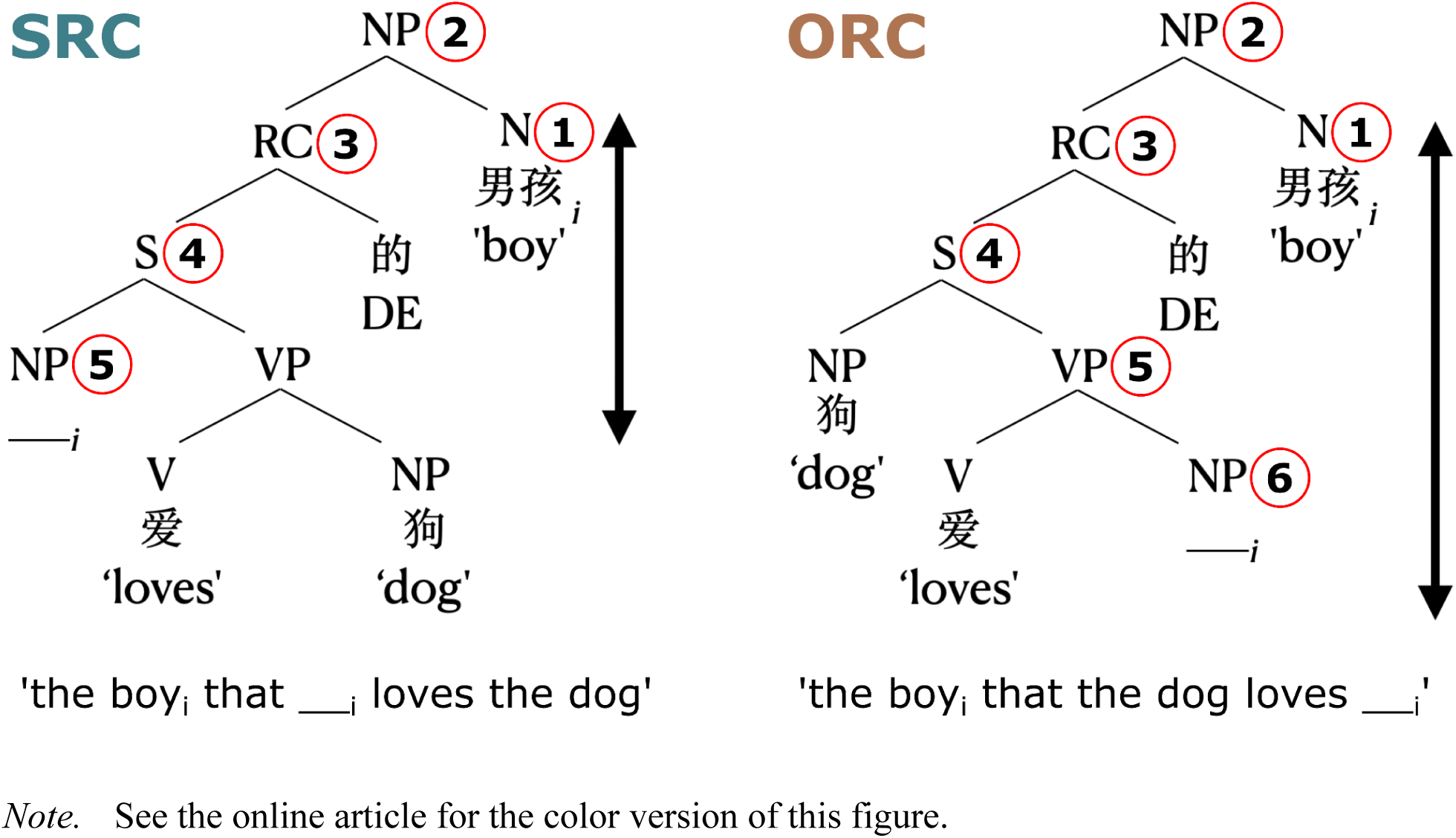
Syntactic Trees for Example Chinese Subject- and Object-Extracted Relative Clauses, Annotated with the Number of Intervening Syntactic Nodes that Occur between the Filler and the Gap Site.

In the behavioral data, an interaction term is observed in which ORC processing is facilitated in the object-modification condition. This (partial) result is in line with the Parallel Function Hypothesis of Sheldon (1974), which predicts a processing advantage for object-modifying ORCs and subject-modifying SRCs. In this account, RC processing is easier when coreferential NPs have the same grammatical function. In observing this effect in the context of RPVP, it may be the case that, due to the demands of the presentation paradigm, participants are (partially) falling back upon a heuristic, child-like processing strategy (Sheldon, 1974).

### Comparison with Serial Presentation

With its novel stimulus presentation paradigm, the EEG results of the study, here, are challenging to square with EEG results from previous serial-presentation investigations of Chinese RC processing. Some differences can be explained through the presence/absence of temporary ambiguity. Yang et al. (2010) find a P600 for SRCs and an N400 for ORCs at the RC verb, which they interpret as structural reanalysis and meaning reanalysis, respectively.

Additionally interpreted as reanalysis, Bulut et al. (2018) observe a complex N400-P600 for ORCs and Xiong et al. (2019) observe a P600 for object-modifying ORCs at the relativizer. As RPVP eliminates main clause-/RC-reading ambiguity (see Hsiao & Gibson, 2003; Lin & Bever, 2010), we would not expect to observe those effects here.

Interestingly, in previous serial EEG work, both P600-structural/syntactic integration and negativity-working memory/referent resolution results are quite consistent. P600s have been observed for SRCs at the relativizer (Packard et al., 2010) and the RC head (Bulut et al., 2018; Packard et al., 2010) in Chinese (but see also a centro-posterior positivity for ORCs at the head noun in Japanese in Ueno & Garnsey, 2008). Negativities for ORCs have been observed at the relativizer (Xiong et al., 2019) and the head noun (Yang et al., 2010) in Chinese, and also between the RC verb and the RC head in Japanese (Ueno & Garnsey, 2008). While working memory effects would not be expected here, we do observe a greater positivity, but for *ORCs* rather than *SRCs*. Additionally, this positivity is not particularly P600-like. Rather than being topographically located over centro-parietal electrodes and occurring between 500-900ms, the positivity observed here is located over right lateral electrodes and comes online ∼400ms post second screen presentation. Participants appear, seemingly due to the particular demands of the task, to be using a different processing strategy or collection of strategies to structurally integrate or referentially resolve the filler with the gap site in RPVP than they would in RSVP.

It is unclear to what degree parallel presentation of sentences (or sentence fragments) necessitates parallel processing. In our study, sentence fragments were displayed for 166ms on average, which is likely too quick for participants to plan and execute a saccade to read the stimuli serially. Nonetheless, it is possible that the processes involved are serial, or that participants in some sense serially process the sentence in visual memory. The kinds of mechanisms deployed in processing sentences displayed so briefly are still under scrutiny (e.g., Dunagan et al., 2024; Flower & Pylkkänen, 2024; Wen et al., 2019). The key point of our study is that even when all elements of the RC dependency are presented within the fovea and parafovea in short sentence fragments that minimize the amount of word-by-word memory computations, an SRC advantage still is observed in behavior and distinct neural signals for SRCs and ORCs are observed. Insofar as parallel presentation necessitates non-serial processes, this is evidence that the SRC advantage does not uniquely derive from the memory computations needed to serially process a RC. However, because of the complexity of our stimuli, we had to break the sentences into two screens which introduced other complications. Our design necessitated some seriality, since the subject NP is processed before the object NP in all stimuli. Additionally, the main verb is included in the first screen for the object-modifying stimuli RC but in the second screen for subject-modifying RC stimuli. If interpreting the main verb and retrieving and reactivating the verb’s arguments from memory is a component of the SRC vs.

ORC difference, then the fact that the verbs do not appear with the same onset time may introduce other complications in interpreting the SRC vs. ORC difference in our study. However, we again point out that main effects of ±Object-Modifying are not of particular theoretical interest, and no interaction effects were observed in the EEG data, suggesting that this had minimal impact on the reported results.

### Conclusion

Here, we presented a novel EEG study examining SRC and ORC processing in Chinese. Unlike prior studies, we used an adapted version of the RPVP paradigm in which the key elements of the RC dependency were presented to the visual field in parallel, mitigating the working memory demands imposed by serial presentation experiments. Despite this novel presentation scheme, we still find an SRC advantage, and distinct EEG signatures for SRC and ORC structures. Thus, the processing difficulty of ORCs vs. SRCs cannot reduce to their different demands on working memory resources as a sentence unfolds word-by-word in speech or longer reading. Instead, we suggest that these effects are more likely to reflect aspects of processing and representing the two grammatical structures. Interestingly, we also observe an interaction effect, in which the SRC advantage is mitigated for object-modifying relative clauses, suggesting that RC processing is partially shaped by the role that the head noun plays in the main clause (Sheldon, 1974).

## Notes

This work was partially supported by UGA Seed funding awarded to Dustin A. Chacón, a UGA Franklin Department SPARK Award awarded to Jill McLendon, NSF Grant 2335767 awarded to Dustin A. Chacón, and NSF Grant 1903783 awarded to John Hale.

https://osf.io/ub3x9/

### Competing Interest Statement

The authors have declared no competing interest.

